## Supplement for "Repurposing the mammalian RNA-binding protein Musashi-1 as an allosteric translation repressor in bacteria"

Roswitha Dolcemascolo, Maria Heras-Hernandez, Lucas Goiriz, Roser Montagud-Martinez, Alejandro Requena-Menendez, Raul Ruiz, Anna Perez-Rafols, R. Anahi Higuera-Rodriguez, Guillermo Perez-Ropero, Wim F. Vranken, Tommaso Martelli, Wolfgang Kaiser, Jos Buijs, and Guillermo Rodrigo

#### Contents

**Supplementary Figures** . . . . . . . . . . . . . . . . . . . . . . . . . . . . . . . **S2**

**Supplementary Tables** . . . . . . . . . . . . . . . . . . . . . . . . . . . . . . . **S9**

**Supplementary Notes** . . . . . . . . . . . . . . . . . . . . . . . . . . . . . . . **S13**

**Supplementary References** . . . . . . . . . . . . . . . . . . . . . . . . . . . . **S18**

### Supplementary Figures

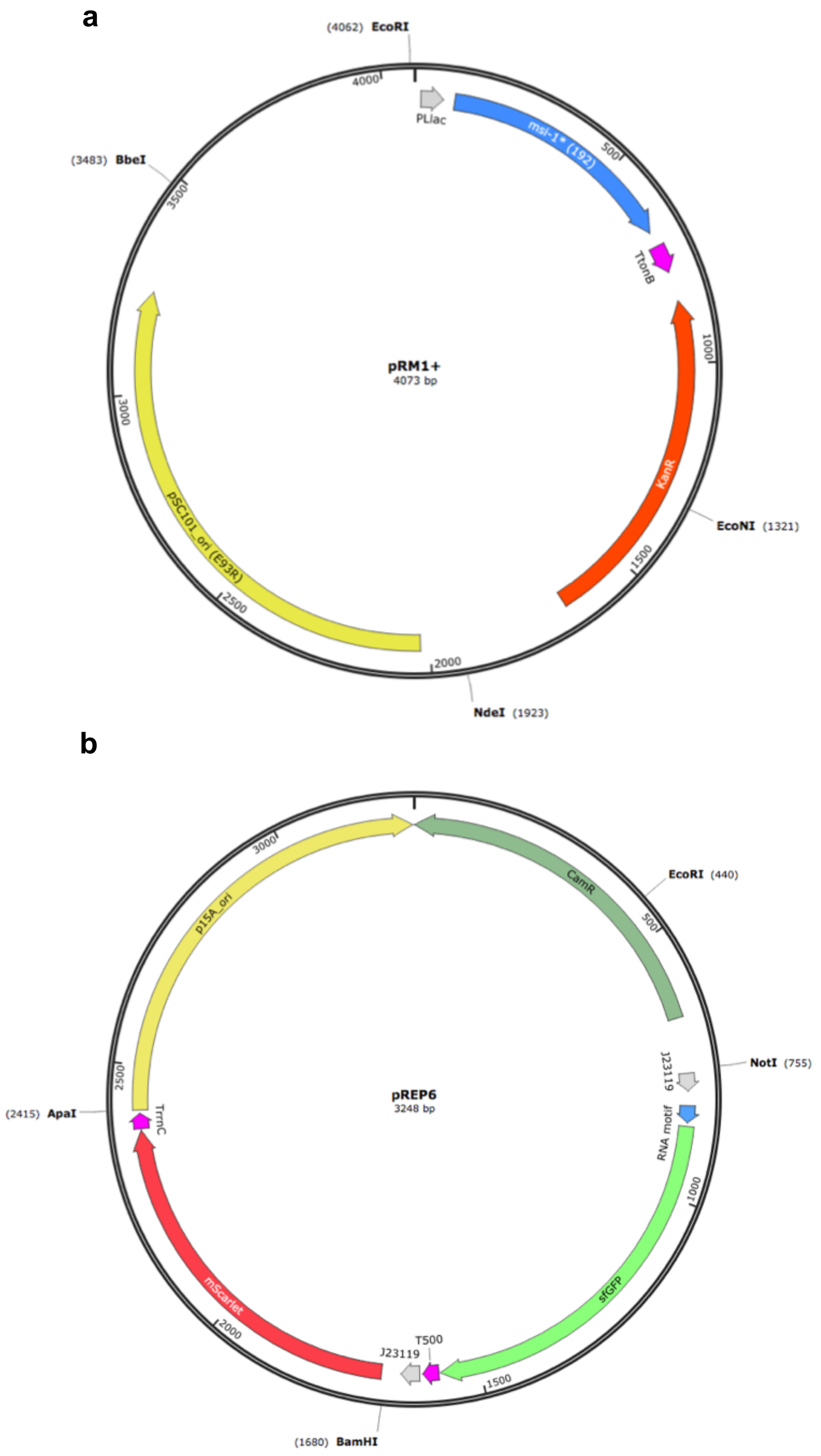


**Supplementary Figure 1:** Maps of the plasmids used to implement the synthetic gene circuit in which MSI-1* represses the translation of sfGFP. a) Map of pRM1+ to express the MSI-1* protein from a PLlac promoter, induced with lactose or IPTG. b) Map of pREP6 to express the reporter sfGFP protein from a constitutive promoter (J23119), harboring a suitable RNA motif in the leader region for translation regulation.

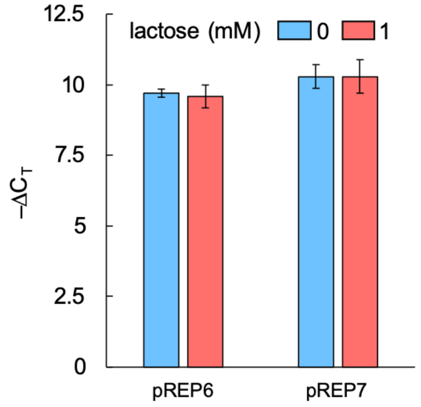


**Supplementary Figure 2:** RT-qPCR results for the mRNA level of *sfGFP* (system implemented with pRM1+ and pREP6 or pREP7, induction with 1 mM lactose), showing no significant change (Welch's $t$-test, two-tailed $P > 0.05$). Error bars correspond to standard deviations ($n = 3$). The oligonucleotide sequences used to amplify the housekeeping gene (in this case, b3500) were TGAGCATCTGGATTACAGCAAC (forward) and CGCGGTGAAAGAGGATTTATAC (reverse) and the probe was TAMRA-TCCGCCGATTGGTACTGTTGGTTT-Q, where the quencher Q was Iowa Black FQ. The oligonucleotide sequences used to amplify *sfGFP* were GTTCAGTGCTTTGCTCGTTATC (forward) and GTACGTGCCGTCATCCTTAAA (reverse) and the probe was FAM-AGCATGACTTCTTCAAGTCCGCCA-Q.

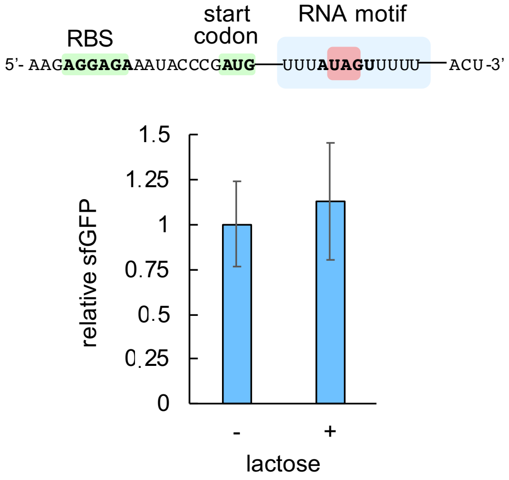


**Supplementary Figure 3:** Characterization of the system response with lactose using pREP4 as a reporter plasmid (induction with 1 mM lactose), showing no significant regulation (Welch's $t$-test, two-tailed $P > 0.05$). Error bars correspond to standard deviations ($n = 3$). On the top, sequence of the leader RNA of *sfGFP* in this case. While MSI-1* can bind to this sequence *in vitro*, these data show that an efficient interaction *in vivo* requires two copies of the consensus recognition sequence for the two RRMs.

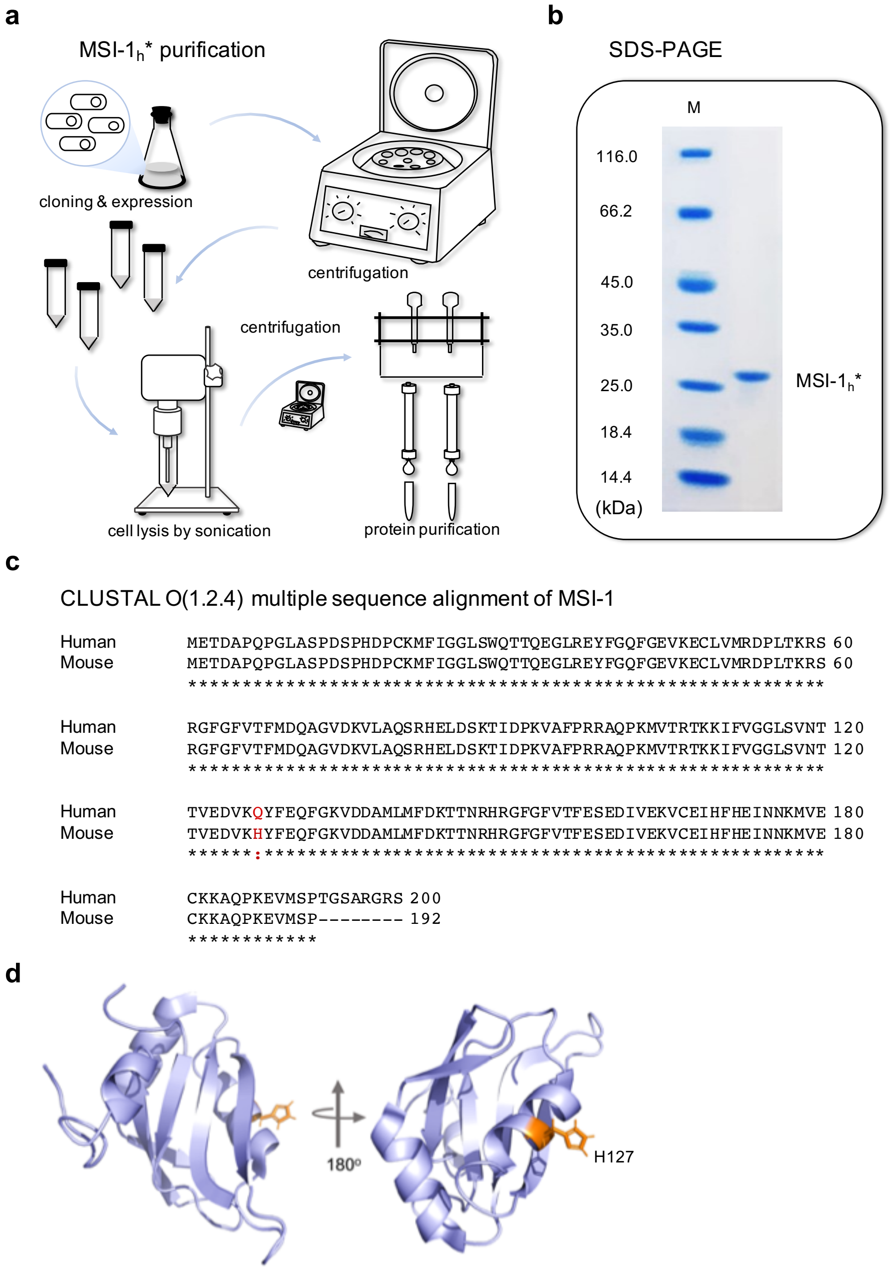


**Supplementary Figure 4:** a) Schematics of the experimental procedure for MSI-$1_h$* purification. *E. coli* cells expressing MSI-$1_h$* were lysed, nucleic acids were removed, and the protein was purified by ion exchange chromatography. b) Gel electrophoretic assay to confirm the presence of MSI-$1_h$*. In particular, this was a sodium dodecyl sulfate polyacrylamide gel electrophoresis (SDS-PAGE). M, molecular marker (Pierce unstained protein MW marker, 14.4-116 kDa, Thermo). c) Sequence comparison between the human and the mouse MSI-1 proteins. The only different residue is located in the first helix of the RRM2. It is residue 127, a histidine in mice and glutamine in humans. This is not expected to make a significant difference in structure or function. The protein that was purified is the human version (denoted by MSI-$1_h$*), while the protein that implements the regulatory circuit is the mouse version (denoted by MSI-1*). d) Three-dimensional structural schematic of the RRM2 of mouse MSI-1.

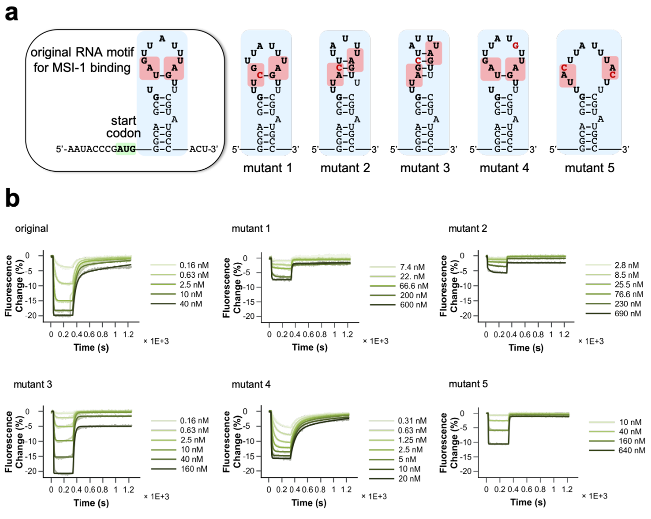


c

| RNA motif ligand | replicate | $k_{ON}$ ($M^{-1}s^{-1}$) | $k_{OFF}$ ($s^{-1}$) | $K_D$ (nM) |
|---|---|---|---|---|
| orig | 1 | (23.600 ± 0.400) × $10^6$ | (10.10 ± 0.10) × $10^{-3}$ | 0.427 ± 0.007 |
| orig | 2 | (17.000 ± 0.300) × $10^6$ | (13.00 ± 0.10) × $10^{-3}$ | 0.760 ± 0.010 |
| orig | 3 | (14.200 ± 0.400) × $10^6$ | (10.40 ± 0.10) × $10^{-3}$ | 0.728 ± 0.010 |
| mut1 | 1 | (22.600 ± 2.400) × $10^6$ | (33.10 ± 1.40) × $10^{-3}$ | 1.470 ± 0.170 |
| mut1 | 2 | (25.100 ± 3.500) × $10^6$ | (22.10 ± 0.90) × $10^{-3}$ | 0.880 ± 0.100 |
| mut1 | 3 | (30.600 ± 3.900) × $10^6$ | (90.90 ± 1.40) × $10^{-3}$ | 2.970 ± 0.380 |
| mut2 | 1 | (16.700 ± 2.000) × $10^6$ | (40.80 ± 2.20) × $10^{-3}$ | 2.440 ± 0.320 |
| mut2 | 2 | (12.600 ± 3.300) × $10^6$ | (27.20 ± 2.90) × $10^{-3}$ | 2.160 ± 0.600 |
| mut2 | 3 | (5.700 ± 1.710) × $10^6$ | (54.40 ± 3.20) × $10^{-3}$ | 9.550 ± 2.930 |
| mut3 | 1 | (5.700 ± 0.210) × $10^6$ | (45.40 ± 0.50) × $10^{-3}$ | 7.970 ± 0.300 |
| mut3 | 2 | (9.000 ± 0.200) × $10^6$ | (31.50 ± 0.20) × $10^{-3}$ | 3.510 ± 0.100 |
| mut3 | 3 | (6.950 ± 0.530) × $10^6$ | (31.60 ± 1.00) × $10^{-3}$ | 4.540 ± 0.370 |
| mut4 | 1 | (14.600 ± 0.200) × $10^6$ | (8.80 ± 0.04) × $10^{-3}$ | 0.600 ± 0.008 |
| mut4 | 2 | (17.000 ± 0.400) × $10^6$ | (12.30 ± 0.10) × $10^{-3}$ | 0.723 ± 0.018 |
| mut4 | 3 | (14.200 ± 0.300) × $10^6$ | (7.96 ± 0.07) × $10^{-3}$ | 0.550 ± 0.014 |
| mut5 | 1 | (0.596 ± 0.060) × $10^6$ | (60.70 ± 2.50) × $10^{-3}$ | 102.000 ± 11.000 |
| mut5 | 2 | (1.540 ± 0.190) × $10^6$ | (87.60 ± 3.60) × $10^{-3}$ | 57.000 ± 7.300 |
| mut5 | 3 | (1.760 ± 0.180) × $10^6$ | (202.00 ± 5.00) × $10^{-3}$ | 115.000 ± 12.000 |

**Supplementary Figure 5:** Characterization of different mutant RNA motifs in terms of binding kinetics against the MSI-$1_h$* protein. a) Predicted secondary structures of original and mutant RNA motifs; mutations are marked in red. b) Binding and unbinding kinetic curves for the different RNA sequences (representative samples). The kinetic constants were extracted from mono-exponential model fits. For the original, mutant 1, mutant 2, and mutant 4 RNA motifs, a bi-exponential model was also explored to describe the binding kinetics, reflecting two types of interactions, although with little improvement. c) Inferred kinetic constants ($k_{\mathrm{ON}}$, $k_{\mathrm{OFF}}$) and the resulting dissociation constant ($K_{\mathrm{D}}$) for each sequence and replicate.

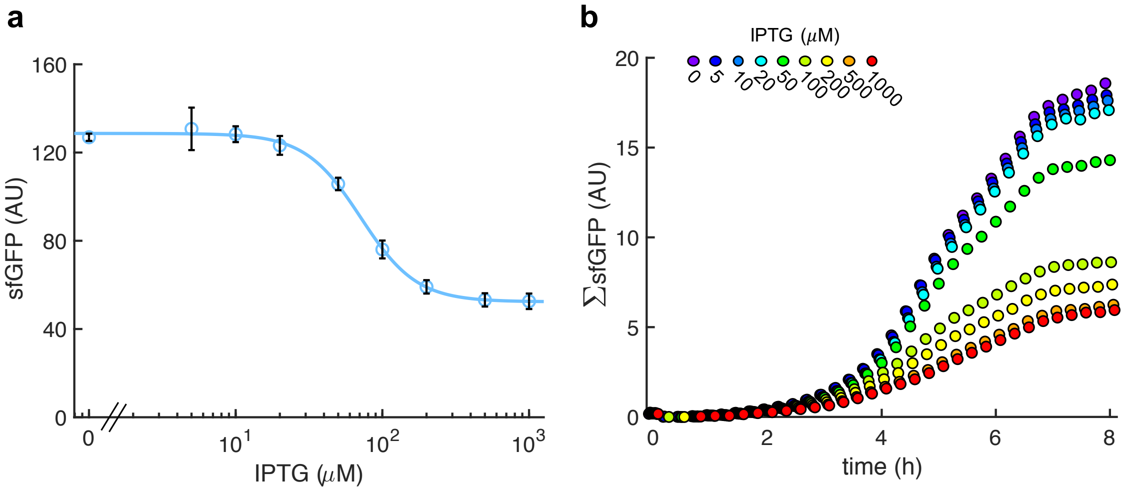


**Supplementary Figure 6:** Characterization of the system response with IPTG (implemented with pRM1+ and pREP6). a) Dose-response curve. Error bars correspond to standard deviations ($n$ = 3). b) Time-course response for different inducer concentrations (average of 4 clones). The fluorescence of the whole population is represented (ΣsfGFP).

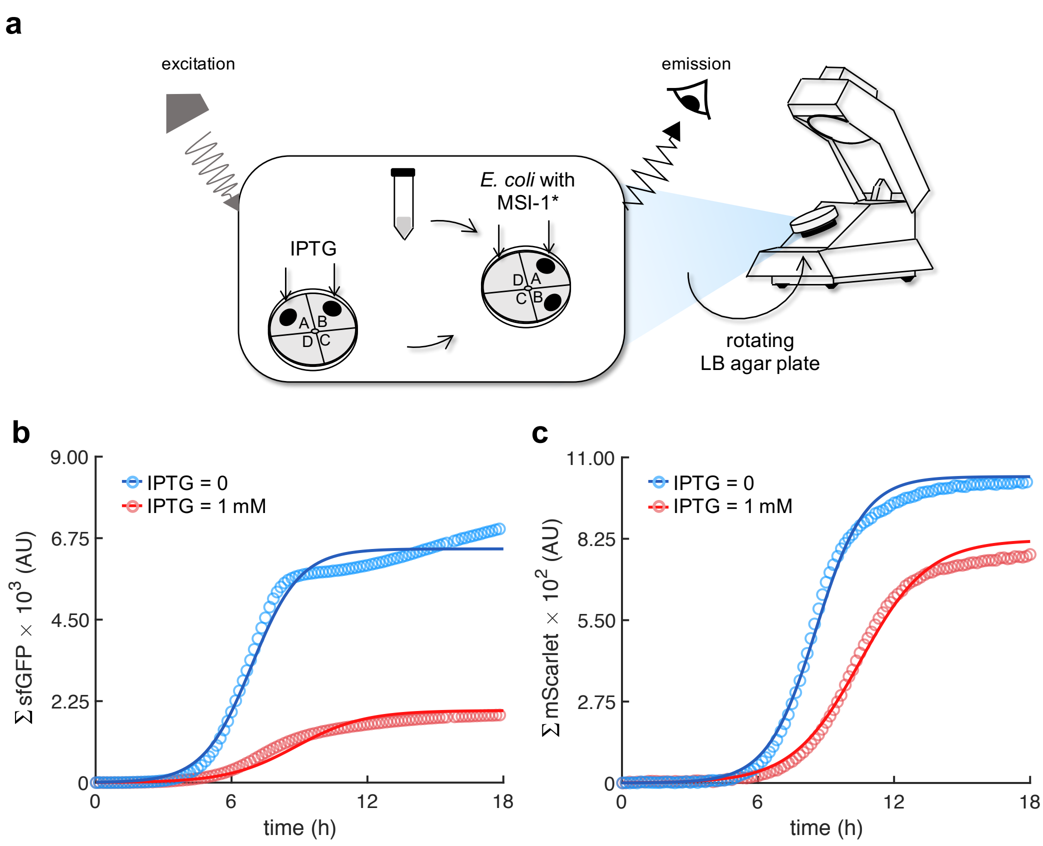


**Supplementary Figure 7:** a) Schematics of the LigandTracer technology repurposed for characterizing bacterial cells expressing fluorescent proteins (implemented with pRM1+ and pREP6). b) Real-time green fluorescence of the whole population (ΣsfGFP) upon induction with IPTG. c) Real-time red fluorescence (ΣmScarlet). Points correspond to the experimental data, while solid lines come from an adjusted mathematical model.

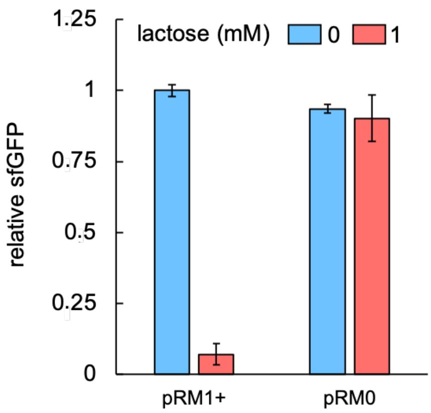


**Supplementary Figure 8:** PLlac promoter tightly controls MSI-1* expression. Characterization of the response with lactose using pREP7 as a reporter plasmid (induction with 1 mM lactose), showing similar fluorescence values in absence of lactose with and without *msi-1**, as well as no significant fluorescence change with lactose in absence of *msi-1**. Error bars correspond to standard deviations ($n = 3$). pRM0 is a void plasmid that does not harbor the *msi-1** genetic cassette.

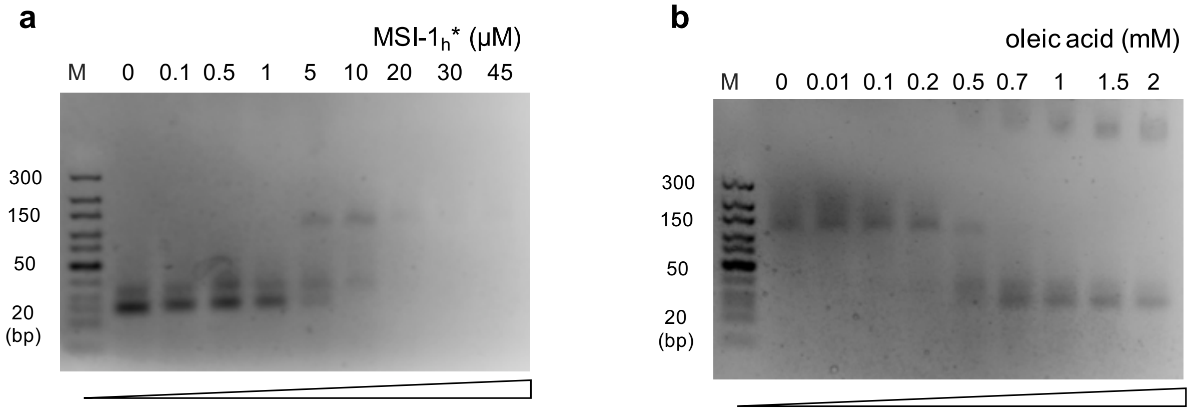


**Supplementary Figure 9:** Gel electrophoresis mobility shift assays to test the MSI-1*-RNA and the MSI-1*-oleic acid interactions (nucleic acid-stained gels). a) Interaction of MSI-1* with the RNA motif. RNA added at 11 $\mu$M. b) Interaction of MSI-1* with oleic acid. RNA added at 11 $\mu$M and MSI-1* at 45 $\mu$M. M, molecular marker (GeneRuler ultra-low range DNA ladder, 10-300 bp, Thermo).

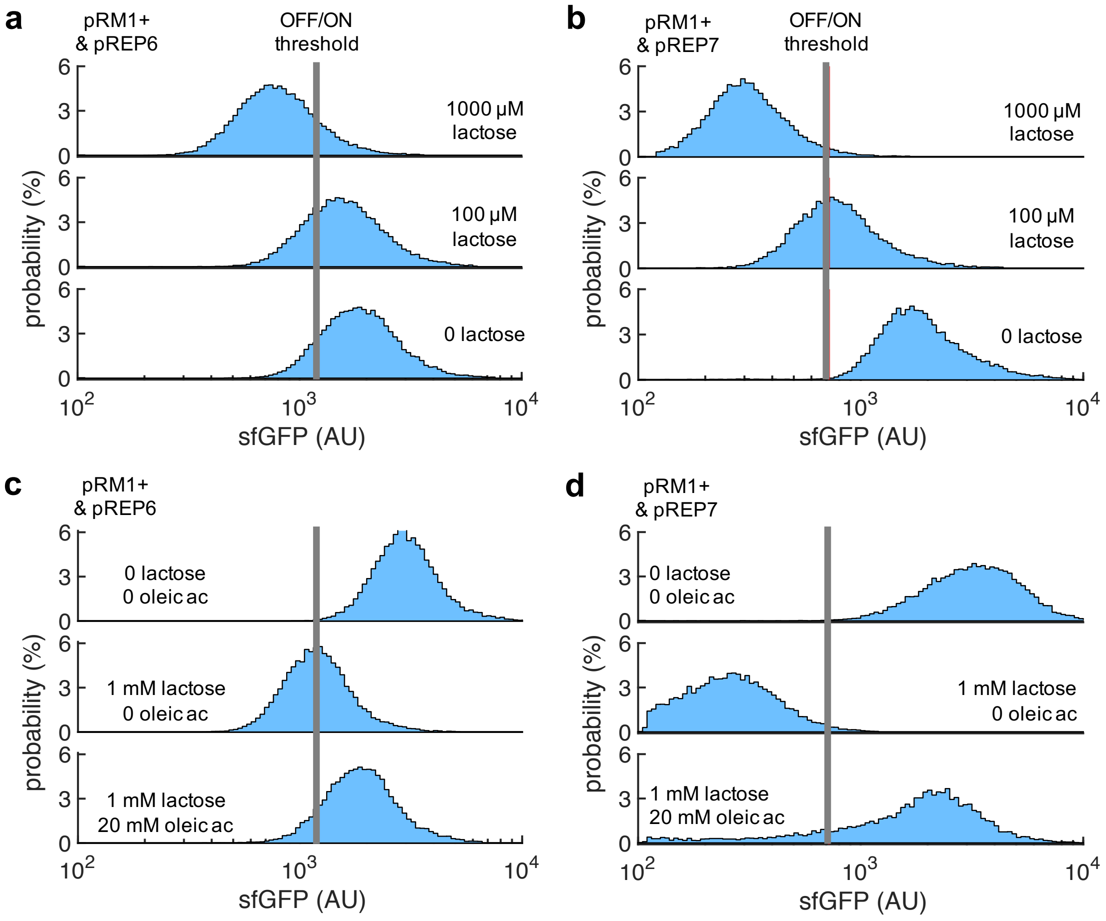


**Supplementary Figure 10:** 2D visualization of probability-based histograms of sfGFP expression from single-cell data. a) System implemented with pRM1+ and pREP6 in response to lactose. This corresponds to Fig. 1f. b) System implemented with pRM1+ and pREP7 in response to lactose. This corresponds to Fig. 4c. c) System implemented with pRM1+ and pREP6 in response to lactose and oleic acid. This corresponds to Fig. 5c. d) System implemented with pRM1+ and pREP7 in response to lactose and oleic acid. This corresponds to Fig. 5d.

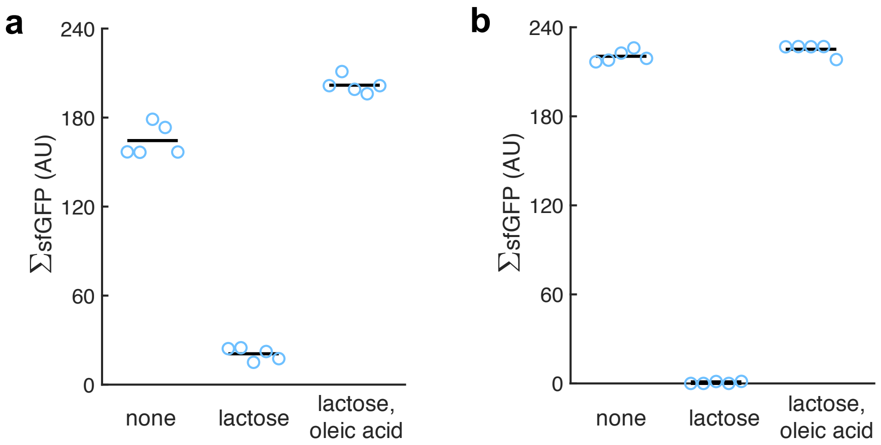


**Supplementary Figure 11:** Quantification of the green fluorescence of the colonies (denoted by ΣsfGFP as it is from populations; $n = 5$). a) From images of *E. coli* colonies harboring pRM1+ and pREP6. b) From images of *E. coli* colonies harboring pRM1+ and pREP7. Oleic acid produced a statistically significant response in both cases (Welch's $t$-test, two-tailed $P < 10^{-4}$).

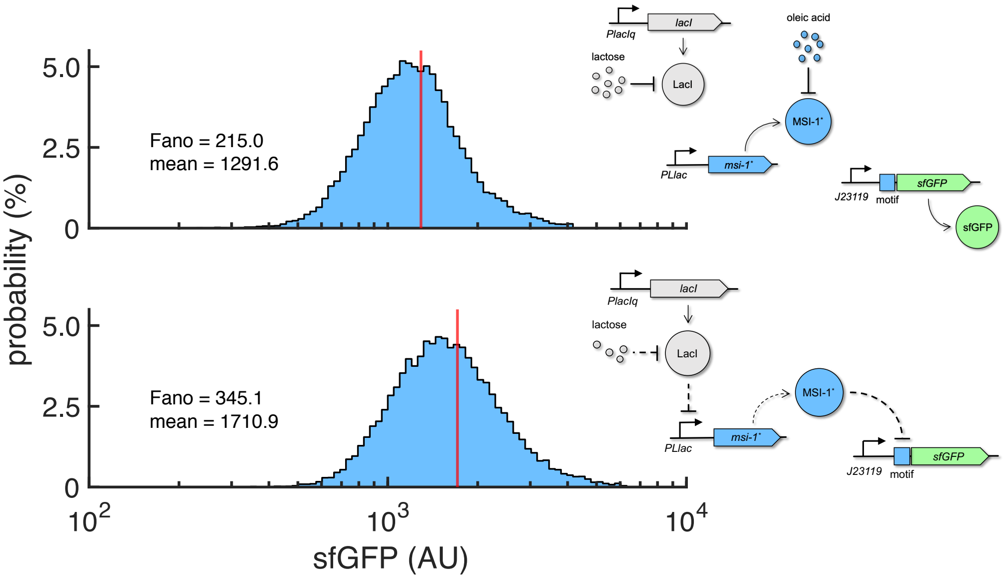


**Supplementary Figure 12:** Probability-based histograms of sfGFP expression from single-cell data for different inducer concentrations (1 mM lactose + 20 mM oleic acid on the top, 0.1 mM lactose on the bottom). The mean expression and Fano factor are shown. Implementation with pRM1+ and pREP6. Schematics of the working modes of the synthetic gene circuit are shown.

### Supplementary Tables

**Supplementary Table 1:** List of plasmids used in this work.

| Name | Insert features | Backbone features | Reference |
|---|---|---|---|
| pRM1+ | PLlac:*msi-1** | KanR, pSC101(E93R) ori | This work |
| pRM0 | void | KanR, pSC101(E93R) ori | This work |
| pRKFR2 | PLlac:*eBFP2* | KanR, pSC101(E93K) ori | [S1] |
| pREP6 | J23119:*sfGFP* (with RNA motif for MSI-1* binding) | CamR, p15A ori | This work |
| pREP6-mut1 | J23119:*sfGFP* (with mutated RNA motif for MSI-1* binding) | CamR, p15A ori | This work |
| pREP6-mut2 | J23119:*sfGFP* (with mutated RNA motif for MSI-1* binding) | CamR, p15A ori | This work |
| pREP6-mut3 | J23119:*sfGFP* (with mutated RNA motif for MSI-1* binding) | CamR, p15A ori | This work |
| pREP6-mut4 | J23119:*sfGFP* (with mutated RNA motif for MSI-1* binding) | CamR, p15A ori | This work |
| pREP6-mut5 | J23119:*sfGFP* (with mutated RNA motif for MSI-1* binding) | CamR, p15A ori | This work |
| pREP7 | J23119:*sfGFP* (with RNA motif for MSI-1* binding and consensus sequences within RBS) | CamR, p15A ori | This work |
| pREP4 | J23119:*sfGFP* (with minimal RNA motif for MSI-1* binding) | CamR, p15A ori | This work |
| pREP4b | J23119:*sfGFP* (with less structured RNA motif for MSI-1* binding) | CamR, p15A ori | This work |
| pREP4b3x | J23119:*sfGFP* (with 3x less structured RNA motifs for MSI-1* binding) | CamR, p15A ori | This work |

| | | | |
|---|---|---|---|
| pGio | T7p:*msi-1$_h$** | KanR, pUC ori | This work |
| pREP6$\alpha$ | PLtet:*sfGFP-mScarlet* (with RNA motif for MSI-1* binding in front of *sfGFP*) | CamR, p15A ori | This work |
| pREP7$\alpha$ | PLtet:*sfGFP-mScarlet* (with RNA motif for MSI-1* binding and consensus sequences within RBS in front of *sfGFP*) | CamR, p15A ori | This work |

**Supplementary Table 2:** Nucleotide sequences of the elements used to implement our synthetic gene circuits.

| **Name** | **Sequence** |
|---|---|
| PLlac | AATTGTGAGCGGATAACAATTGACATTGTGAGCGGATAACAAGATACTGAGCAC |
| *msi-1** (codon optimized to *E. coli* from *M. musculus*) | ATGGAAACGGACGCCCCGCAGCCGGGACTGGCCTCTCCTGACTCTCCTCACGACCCA<br>TGCAAGATGTTTATTGGTGGACTTTCTTGGCAGACTACTCAGGAGGGTCTTCGTGAA<br>TACTTCGGTCAATTTGGCGAAGTGAAAGAGTGTCTTGTGATGCGCGATCCTTTAACC<br>AAGCGTAGTCGCGGATTTGGCTTCGTCACGTTCATGGACCAGGCAGGCGTGGATAAG<br>GTGCTGGCGCAGAGTCGTCACGAATTAGATTCAAAAACGATTGACCCCAAAGTGGCG<br>TTCCCACGTCGCGCCCAACCTAAAATGGTTACTCGTACCAAAAAGATTTTCGTAGGA<br>GGCTTATCCGTAAATACCACGGTAGAAGATGTAAAGCATTACTTCGAACAGTTTGGA<br>AAGGTGGATGATGCAATGCTTATGTTTGATAAGACCACAAACCGTCATCGTGGATTC<br>GGCTTTGTGACCTTTGAATCGGAGGATATCGTTGAGAAGGTCTGCGAAATCCACTTT<br>CATGAAATTAATAACAAAATGGTTGAGTGTAAGAAGGCGCAACCGAAAGAAGTCATG<br>TCTCCTTAA |
| J23119 | TTGACAGCTAGCTCAGTCCTAGGTATAATGCTAGC |
| PLtet | TCCCTATCAGTGATAGAGATTGACATCCCTATCAGTGATAGAGATACTGAGCAC |
| *sfGFP* (RNA motif underlined) | ATGGGCAGCGTTAGTTATTTAGTTCGTATGCCAACTAGTCGTAAAGGCGAAGAGCTG<br>TTCACTGGTGTCGTCCCTATTCTGGTGGAACTGGATGGTGATGTCAACGGTCATAAG<br>TTTTCCGTGCGTGGCGAGGGTGAAGGTGACGCAACTAATGGTAAACTGACGCTGAAG<br>TTCATCTGTACTACTGGTAAACTGCCGGTACCTTGGCCGACTCTGGTAACGACGCTG<br>ACTTATGGTGTTCAGTGCTTTGCTCGTTATCCGGACCATATGAAGCAGCATGACTTC<br>TTCAAGTCCGCCATGCCGGAAGGCTATGTGCAGGAACGCACGATTTCCTTTAAGGAT<br>GACGGCACGTACAAAACGCGTGCGGAAGTGAAATTTGAAGGCGATACCCTGGTAAAC<br>CGCATTGAGCTGAAAGGCATTGACTTTAAAGAAGACGGCAATATCCTGGGCCATAAG<br>CTGGAATACAATTTTAACAGCCACAATGTTTACATCACCGCCGATAAACAAAAAAAT<br>GGCATTAAAGCGAATTTTAAAATTCGCCACAACGTGGAGGATGGCAGCGTGCAGCTG<br>GCTGATCACTACCAGCAAAACACTCCAATCGGTGATGGTCCTGTTCTGCTGCCAGAC<br>AATCACTATCTGAGCACGCAAAGCGTTCTGTCTAAAGATCCGAACGAGAAACGCGAT<br>CATATGGTTCTGCTGGAGTTCGTAACCGCAGCGGGCATCACGCATGGTATGGATGAA<br>CTGTACAAATAA |
| RNA motif mutant 1 (A>C substitution) | GGCAGCGTTCGTTATTTAGTTCGTATGCC |
| RNA motif mutant 2 (G>C substitution) | GGCAGCGTTACTTATTTAGTTCGTATGCC |
| RNA motif mutant 3 (T>C substitution) | GGCAGCGTTAGCTATTTAGTTCGTATGCC |
| RNA motif mutant 4 (G insertion, a nucleotide was deleted downstream to be in frame) | GGCAGCGTTAGTTATGTTAGTTCGTATGCC |

| | |
|---|---|
| RNA motif mutant 5 (two G>C substitutions) | `GGCAGCGTTACTTATTTACTTCGTATGCC` |
| Consensus sequences within RBS | `ATCATGTGTTTAGTTAGGAGATTTAGTTA` |
| Minimal RNA motif | `TTTATAGTTT` |
| Less structured RNA motif | `GGCGTTAGTTATTTAGTTCGCC` |
| 3x less structured RNA motifs | `GGCGTTAGTTATTTAGTTCGCCGACGCGTTAGTTTATTTAGTTCGCGATATCCGGTT`<br>`AGTTATTTAGTTACGG` |
| *mScarlet* | `ATGGGATCCGTGAGCAAGGGCGAGGCAGTGATCAAGGAGTTCATGCGGTTCAAGGTG`<br>`CACATGGAGGGCTCCATGAACGGCCACGAGTTCGAGATCGAGGGCGAGGGCGAGGGC`<br>`CGCCCCTACGAGGGCACCCAGACCGCCAAGCTGAAGGTGACCAAGGGTGGCCCCCTG`<br>`CCCTTCTCCTGGGACATCCTGTCCCCTCAGTTCATGTACGGCTCCAGGGCCTTCATC`<br>`AAGCACCCCGCCGACATCCCCGACTACTATAAGCAGTCCTTCCCCGAGGGCTTCAAG`<br>`TGGGAGCGCGTGATGAACTTCGAGGACGGCGGCGCCGTGACCGTGACCCAGGACACC`<br>`TCCCTGGAGGACGGCACCCTGATCTACAAGGTGAAGCTCCGCGGCACCAACTTCCCT`<br>`CCTGACGGCCCCGTAATGCAGAAGAAGACAATGGGCTGGGAAGCGTCCACCGAGCGG`<br>`TTGTACCCCGAGGACGGCGTGCTGAAGGGCGACATTAAGATGGCCCTGCGCCTGAAG`<br>`GACGGCGGCCGTTACCTGGCGGACTTCAAGACCACCTACAAGGCCAAGAAGCCCGTG`<br>`CAGATGCCCGGCGCCTACAACGTCGACCGCAAGTTGGACATCACCTCCCACAACGAG`<br>`GACTACACCGTGGTGGAACAGTACGAACGCTCCGAGGGCCGCCACTCCACCGGCGGC`<br>`ATGGACGAGCTGTACAAGTAA` |
| *eBFP2* | `ATGGTGAGCAAGGGCGAGGAGCTGTTCACCGGGGTGGTGCCCATCCTGGTCGAGCTG`<br>`GACGGCGACGTAAACGGCCACAAGTTCAGCGTGAGGGGCGAGGGCGAGGGCGATGCC`<br>`ACCAACGGCAAGCTGACCCTGAAGTTCATCTGCACCACCGGCAAGCTGCCCGTGCCC`<br>`TGGCCCACCCTCGTGACCACCCTGAGCCACGGCGTGCAGTGCTTCGCCCGCTACCCC`<br>`GACCACATGAAGCAGCACGACTTCTTCAAGTCCGCCATGCCCGAAGGCTACGTCCAG`<br>`GAGCGCACCATCTTCTTCAAGGACGACGGCACCTACAAGACCCGCGCCGAGGTGAAG`<br>`TTCGAGGGCGACACCCTGGTGAACCGCATCGAGCTGAAGGGCGTCGACTTCAAGGAG`<br>`GACGGCAACATCCTGGGGCACAAGCTGGAGTACAACTTCAACAGCCACAACATCTAT`<br>`ATCATGGCCGTCAAGCAGAAGAACGGCATCAAGGTGAACTTCAAGATCCGCCACAAC`<br>`GTGGAGGACGGCAGCGTGCAGCTCGCCGACCACTACCAGCAGAACACCCCCATCGGC`<br>`GACGGCCCCGTGCTGCTGCCCGACAGCCACTACCTGAGCACCCAGTCCGTGCTGAGC`<br>`AAAGACCCCAACGAGAAGCGCGATCACATGGTCCTGCTGGAGTTCCGCACCGCCGCC`<br>`GGGATCACTCTCGGCATGGACGAGCTGTACAAG` |

### Supplementary Notes

#### Supplementary Note 1

The repression of sfGFP as a function of lactose is modelled by the following Hill equation [S2]

$$[\text{sfGFP}] = \frac{A_1}{1 + \left(\frac{[\text{Lactose}]}{K_1}\right)^{n_1}} + B_1 \ ,$$

where $K_1$ is the regulatory coefficient, $n_1$ the Hill coefficient, $A_1 + B_1$ the maximal expression level, and $B_1$ the basal expression level at full repression. In the case of sfGFP, its concentration is given by the normalized green fluorescence signal in arbitrary units (AU). The adjusted parameter values are $A_1 = 90.7$ AU, $B_1 = 62.1$ AU, $K_1 = 99.1$ $\mu$M, and $n_1 = 1.70$ (Fig. 1c).

We used the very same equation to model the dynamic response of the system implemented with pREP7. In this case, the adjusted parameter values are $A_1 = 289.9$ AU, $B_1 = 39.4$ AU, $K_1 = 85.7$ $\mu$M, and $n_1 = 4.45$ (Fig. 4b).

Also, the following Hill equation models the repression of sfGFP by IPTG

$$[\text{sfGFP}] = \frac{A_2}{1 + \left(\frac{[\text{IPTG}]}{K_2}\right)^{n_2}} + B_2 \ .$$

The adjusted parameter values are $A_2 = 76.4$ AU, $B_2 = 52.3$ AU, $K_2 = 71.3$ $\mu$M, and $n_2 = 2.28$ (Suppl. Fig. 5a).

In addition, the activation of eBFP2, proxy of MSI-1*, as a function of lactose is modelled by the following Hill equation

$$[\text{MSI-1*}] \propto [\text{eBFP2}] = \frac{A_3 \left(\frac{[\text{Lactose}]}{K_3}\right)^{n_3}}{1 + \left(\frac{[\text{Lactose}]}{K_3}\right)^{n_3}} + B_3 \ ,$$

where $K_3$ is the regulatory coefficient, $n_3$ the Hill coefficient, $A_3 + B_3$ the maximal expression level, and $B_3$ the basal expression level with no activation. In the case of eBFP2, its concentration is given by the normalized blue fluorescence signal in AU. The adjusted parameter values are $A_3 = 23.1$ AU, $B_3 = 1.88$ AU, $K_3 = 359$ $\mu$M, and $n_3 = 2.81$ (Fig. 1d).

Finally, the following Michaelis equation (a particular case of the Hill equation when there is no cooperativity)

$$[\text{sfGFP}] = \frac{A_4}{1 + \frac{[\text{eBFP2}]}{K_4}}$$

defines the engineered regulation between MSI-1* (given by eBFP2) and sfGFP. Here, no basal expression level is considered. The adjusted parameter values are $A_4 = 165$ AU and $K_4 = 10.2$ AU (Fig. 1d).

**Supplementary Note 2**

The fold change in protein expression can be calculated from the fundamental parameters that model the regulatory system, such as the association rate of the regulator to the nucleic acid ($k_{\mathrm{ON}}$), the dissociation rate ($k_{\mathrm{ON}}$), the concentration of the regulator in the cell ($R$), and the degradation rate of the nucleic acid ($\delta$). If we denote by $A_0$ the concentration of free nucleic acid, by $A_R$ the concentration of nucleic acid with the regulator bound, and by $P$ the concentration of the regulated protein, we can write

$$\begin{aligned}\frac{dA_0}{dt} &= \alpha - k_{\mathrm{ON}} R A_0 + k_{\mathrm{OFF}} A_R - \delta A_0 \\ \frac{dA_R}{dt} &= k_{\mathrm{ON}} R A_0 - k_{\mathrm{OFF}} A_R - \delta A_R \\ \frac{dP}{dt} &= \beta A_0 + \varepsilon \beta A_R - \mu P \ ,\end{aligned}$$

where $\alpha$ is the synthesis rate of the nucleic acid, $\beta$ the synthesis rate of the protein, and $\varepsilon$ the leakage fraction of protein synthesis when the regulator is bound. Note that in steady state $A_{0\infty} + A_{R\infty} = \dfrac{\alpha}{\delta}$.

If $R = 0$, then $P_\infty = \dfrac{\alpha\beta}{\delta\mu}$ (steady state). If $R > 0$, then $P_\infty = \dfrac{\alpha\beta}{\delta\mu}\left(\dfrac{\varepsilon k_{\mathrm{ON}} R + k_{\mathrm{OFF}} + \delta}{k_{\mathrm{ON}} R + k_{\mathrm{OFF}} + \delta}\right)$. Therefore, it turns out that

$$\text{fold} = \frac{P_\infty(R=0)}{P_\infty(R>0)} = \frac{k_{\mathrm{ON}} R + k_{\mathrm{OFF}} + \delta}{\varepsilon k_{\mathrm{ON}} R + k_{\mathrm{OFF}} + \delta} \ .$$

Importantly, this model can be applied either to transcription regulation or translation regulation. The main difference is that in the case of transcription, the nucleic acid targeted by the regulator (DNA) is stable (we can model this as $\delta = \mu$, and then set $\delta \simeq 0$ in the fold change equation), while in the case of translation, the nucleic acid targeted by the regulator (mRNA) is unstable ($\delta \gg \mu$).

**Supplementary Note 3**

The system of ordinary differential equations (ODEs) that governs the dynamics of the engineered circuit, considering the intracellular concentrations of mRNAs and proteins [S3], reads

$$
\begin{aligned}
\frac{d\,[\mathrm{mRNA}_{\text{MSI-1*}}]}{dt} &= \alpha_x \left( \frac{\rho_x + \left(\frac{[\text{Lactose}]}{\theta_x}\right)^{n_x}}{1 + \left(\frac{[\text{Lactose}]}{\theta_x}\right)^{n_x}} \right) - \delta\,[\mathrm{mRNA}_{\text{MSI-1*}}] \\
\frac{d\,[\text{MSI-1*}]}{dt} &= \beta_x\,[\mathrm{mRNA}_{\text{MSI-1*}}] - \mu\,[\text{MSI-1*}] \\
\frac{d\,[\mathrm{mRNA}_{\text{sfGFP}}]}{dt} &= \alpha_y - \delta\,[\mathrm{mRNA}_{\text{sfGFP}}] \\
\frac{d\,[\text{sfGFP}]}{dt} &= \beta_y \left( \frac{1}{1 + \frac{[\text{MSI-1*}]}{\theta_y}} \right) [\mathrm{mRNA}_{\text{sfGFP}}] - \mu\,[\text{sfGFP}] \ ,
\end{aligned}
$$

where $\alpha_x$ is the maximal transcription rate of the *msi-1** gene, $\alpha_y$ the maximal transcription rate of the *sfGFP* gene, $\beta_x$ the maximal translation rate of *msi-1**, $\beta_y$ the maximal translation rate of *sfGFP*, $\delta$ the mRNA degradation rate (assumed equal for the *msi-1** and *sfGFP* genes), $\rho_x$ the repression fold of LacI at the transcriptional level, $\theta_x$ the effective dissociation constant between LacI and lactose, $n_x$ the effective binding cooperativity of LacI, $\theta_y$ the effective dissociation constant between MSI-1* and the RNA motif in the *sfGFP* gene, and $\mu$ the bacterial growth rate.

The analytical solution of this system of ODEs can be obtained through the use of the Laplace transform [S4] and reads

$$
\begin{aligned}
[\mathrm{mRNA}_{\text{MSI-1*}}]\,(t) &= \frac{\alpha_x}{\delta} \left( \frac{\rho_x + \left(\frac{[\text{Lactose}]}{\theta_x}\right)^{n_x}}{1 + \left(\frac{[\text{Lactose}]}{\theta_x}\right)^{n_x}} \right) \left(1 - e^{-\delta t}\right) + [\mathrm{mRNA}_{\text{MSI-1*}}]_0\, e^{-\delta t} \\
[\text{MSI-1*}]\,(t) &= \beta_x \int_0^t e^{-\mu(t-\tau)}\,[\mathrm{mRNA}_{\text{MSI-1*}}]\,(\tau) d\tau + [\text{MSI-1*}]_0\, e^{-\mu t} \simeq \\
&\simeq \frac{\beta_x}{\mu}\,[\mathrm{mRNA}_{\text{MSI-1*}}]_\infty \left(1 - e^{-\mu t}\right) + [\text{MSI-1*}]_0\, e^{-\mu t} \\
[\mathrm{mRNA}_{\text{sfGFP}}]\,(t) &= \frac{\alpha_y}{\delta} \left(1 - e^{-\delta t}\right) + [\mathrm{mRNA}_{\text{sfGFP}}]_0\, e^{-\delta t} \\
[\text{sfGFP}]\,(t) &= \beta_y \int_0^t e^{-\mu(t-\tau)} \left( \frac{[\mathrm{mRNA}_{\text{sfGFP}}]\,(\tau)}{1 + \frac{[\text{MSI-1*}](\tau)}{\theta_y}} \right) d\tau + [\text{sfGFP}]_0\, e^{-\mu t} \simeq \\
&\simeq \frac{\alpha_y \beta_y}{\delta} \int_0^t \frac{e^{-\mu(t-\tau)}}{1 + \frac{[\text{MSI-1*}](\tau)}{\theta_y}} d\tau + [\text{sfGFP}]_0\, e^{-\mu t} \ ,
\end{aligned}
$$

where to perform the approximations $\delta \gg \mu$ is considered (quasi-steady state scenario).

Then, in the steady state, we have

$$[\text{mRNA}_{\text{MSI-1*}}]_\infty = \frac{\alpha_x}{\delta}\left(\frac{\rho_x + \left(\frac{[\text{Lactose}]}{\theta_x}\right)^{n_x}}{1 + \left(\frac{[\text{Lactose}]}{\theta_x}\right)^{n_x}}\right)$$

$$[\text{MSI-1*}]_\infty = \frac{\alpha_x\beta_x}{\delta\mu}\left(\frac{\rho_x + \left(\frac{[\text{Lactose}]}{\theta_x}\right)^{n_x}}{1 + \left(\frac{[\text{Lactose}]}{\theta_x}\right)^{n_x}}\right)$$

$$[\text{mRNA}_{\text{sfGFP}}]_\infty = \frac{\alpha_y}{\delta}$$

$$[\text{sfGFP}]_\infty = \frac{\alpha_y\beta_y}{\delta\mu}\left(\frac{1}{1 + \frac{\alpha_x\beta_x}{\delta\mu\theta_y}\left(\frac{\rho_x + \left(\frac{[\text{Lactose}]}{\theta_x}\right)^{n_x}}{1 + \left(\frac{[\text{Lactose}]}{\theta_x}\right)^{n_x}}\right)}\right).$$

From the growth curves, we calculated $\mu = 0.8$ h$^{-1}$. Knowing that in *E. coli* the average half-life of mRNA is 5 min [S5], we set $\delta = 0.14$ min$^{-1}$. Using our experimental data, the adjusted parameter values are $\frac{\alpha_x\beta_x}{\theta_y} = 13$ h$^{-2}$, $\alpha_y\beta_y = 17$ AU/h$^2$, $\rho_x = 0.075$, $\theta_x = 150$ $\mu$M, and $n_x = 1.5$ (Fig. 3e-g).

**Supplementary Note 4**

The number of cells ($N$) in a bacterial culture with time can be described by a logistic function [S6] as

$$N(t) = \frac{N_{\max}}{1 + e^{-\mu(t-\psi)}} \ ,$$

where $N_{\max}$ is the maximal capacity of the medium, $\mu$ the bacterial growth rate, and $\psi$ the delay of the response (or the time at which the culture reaches half of the capacity).

In our experimental system, the constitutive expression of mScarlet may be used to estimate the total number of cells. Indeed, the absolute red fluorescence level ($\Sigma$mScarlet) may be assumed proportional to $N$. Thus, we may write

$$\Sigma\text{mScarlet}(t) = \frac{\Sigma\text{mScarlet}_{\max}}{1 + e^{-\mu(t-\psi)}}$$
$$\Sigma\text{sfGFP}(t) = [\text{sfGFP}]\,(t) \cdot \Sigma\text{mScarlet}(t).$$

Using our experimental data, the adjusted parameter values are $\Sigma\text{mScarlet}_{\max} = 13.9$ AU in the case of no induction, $\Sigma\text{mScarlet}_{\max} = 12.5$ AU when induced with 1 mM lactose, $\mu = 0.8$ $h^{-1}$, and $\psi = 6.5$ h (Fig. 3d). [sfGFP] $(t)$ was calculated as described in the Suppl. Note 3.

With the time-dependent experimental data in solid media (from LigandTracer), the adjusted parameter values are $\mu = 0.0156$ $\text{min}^{-1}$, $\psi = 513$ min, $\Sigma\text{mScarlet}_{\max} = 1035$ AU, and [sfGFP] $= 6.23$ AU in the case of no induction, and $\mu = 0.0111$ $\text{min}^{-1}$, $\psi = 630$ min, $\Sigma\text{mScarlet}_{\max} = 821$ AU, and [sfGFP] $= 2.42$ AU when induced with 1 mM IPTG (Suppl. Fig. 6). In this case, for simplicity, we considered a quasi-steady state scenario, setting constant the sfGFP expression. Moreover, we noticed a delay of about 100 min ($= \nu$) between the mScarlet and sfGFP expressions, so the equation $\Sigma\text{sfGFP}(t) = [\text{sfGFP}] \cdot \Sigma\text{mScarlet}(t + \nu)$ was used instead to fit the data.
